# Advantage of the F(ab)’2 fragment over IgG for RIT and PRIT

**DOI:** 10.1101/2024.10.16.617029

**Authors:** Zainab Tabaja, Maya Diab, Frédérique Nguyen, Catherine Maurel, Frédéric Pecorari, Sylvia Lambot, François Davodeau

## Abstract

Radioimmunotherapy (RIT) has proved clinically effective in the treatment of relapsed or consolidating NHL after chemotherapy. However, it is still at the clinical trial stage for the treatment of solid tumors. The efficacy of the treatment on this type of tumor is limited by hematological toxicity, which restricts the activities injected. Pre-targeted radioimmunotherapy (PRIT) enables radiolabeling to be carried out in vivo at a time when the quantities of antigen-targeted vector reach a maximum level in the tumor, but have largely decayed in the bloodstream, thus limiting hematological toxicity. This therapeutic approach would make it possible to use higher radiopharmaceutical activities than those used in RIT, thereby increasing the doses deposited in the tumor and the therapeutic response.

Despite encouraging preclinical trials, this approach has failed to live up to expectations in terms of clinical applications. In the context of PRIT using the Inverse Electron Demand Diels Alder (IEDDA) reaction as a method for radiolabeling tumor-targeted vectors, a major research effort has been made to improve the quality of in vivo radiolabeling using tetrazine (Tz) and transcycloctene (TCO) variants whose stability and reactivity have been optimized. Most trials in this field have been carried out using antibodies which, thanks to their long biological half-life, enable prolonged irradiation of tumors. Low-molecular-weight vectors such as affinity proteins or antigen-specific antibody fragments (VHH,scFv, F(ab)’) have the advantage of being rapidly eliminated from circulation by renal filtration, but on the other hand, the quantities and residence times of these vectors in the tumor are reduced compared with IgG.

F(ab)’2 antibody fragments have properties intermediate between those of IgG and low-molecular-weight vectors that could be suitable for PRIT approaches, but they have not yet been evaluated in PRIT using IEDDA. In preparation for a preclinical trial of radioimmunotherapy of triple-negative breast cancer in spontaneously affected dogs, we compared the biodistribution of IgG and its F(ab)’2 fragment targeting canine CD138 in a xenogeneic model of canine breast cancer and in a mouse model of colon cancer expressing human ACE antigen. This analysis shows that the use of F(ab)’2 fragments improves the therapeutic index compared with IgG. F(ab)’2 fragments seem particularly well suited to tumor targeting in PRIT using short-lived isotopes such as 211At.

## Introduction

Pre-targeting tumors with specific vectors improves the contrast between tumor and healthy tissue in nuclear imaging and shortens the time between radiopharmaceutical injection and SPECT/CT or PET/CT image acquisition. Unlike tumor targeting using a directly radiolabeled vector in the classical radioimmunotherapy (RIT) approach, pre-targeting limits the quantity of circulating vector, thereby improving tumor-to-healthy organ ratios.

This targeting approach has been of interest in radioimmunotherapy, where it theoretically makes it possible to avoid doses deposited to healthy organs from circulating vectors during the tumor-specific vector accretion phase. In particular, pre-targeted radioimmunotherapy (PRIT) enables optimal irradiation of tumors with short-lived radioactive isotopes such as alpha-emitting Bismuth-213 (²¹³Bi) or Astatine-211 (²¹¹At).

Three pre-targeting approaches have been widely studied: The first is to use vectors with a valence specific to a tumor antigen and a valence directed against a radiolabeled hapten. The second approach involves using the Avidin/Biotin system, where vectors are coupled to streptavidin and radiolabeled biotin ^1^. More recently, the advent of orthogonal biochemistry has enabled the development of a third pre-targeting method that employs complementary click chemistry functions, such as TCO coupled to the tumor-specific vector and radiolabeled Tetrazine injected as a second step ^2 3^. Other *in-vivo* radiolabeling approaches based on natural or synthetic complementary oligonucleotides, such as morpholino nucleic acids ^4^, nucleic acid peptide ^5^ or endonuclease-resistant L-series oligonucleotides, have also been evaluated ^6^.

In all these approaches, the use of low-molecular-weight radiolabeled hapten results in rapid elimination from the bloodstream through renal filtration. Limiting the circulation time of the radiolabeled hapten reduces toxicity to healthy organs, particularly hematological toxicity, which defines the maximum tolerated dose in one-step RIT approaches. However, renal filtration of the radiolabeled hapten can potentially lead to kidney and bladder toxicity, which may become limiting, especially with alpha emitters. This condition affects the allowable injectable activities.

Most murine tumor models used in preclinical studies to evaluate PRIT strategies exhibit strong expression of the targeted tumor antigen ^7 8^. The range of activity required for effective antitumor dose deposition often exceeds the dose limits for acute toxicity. Despite the proof-of-concept demonstrated in these preclinical models, the results of PRIT approaches carried out with antibody in clinical trials have been short of the expectations set by preclinical assays. Consequently, these trials have yet to expand the application of radioimmunotherapy in the clinic for solid or hematological tumors.

We previously demonstrated that targeting CD138 with the B-B4-^131^I antibody on MDA-MB-468 tumors in triple-negative breast cancer xenografts in nude mice allowed for tumor elimination (100 days) in half of the treated mice at the maximum tolerated injected activity (MTD). Additionally, it significantly reduced tumor volume in those mice showing partial response ^9^. Thus, PRIT offers an alternative to one-step RIT, enhancing treatment efficacy in relapsed triple-negative breast cancer by increasing the MTD. Moreover, PRIT enables the consideration of alpha particle-emitting isotopes for treatment.

Unlike the Tag-72 or A33 antigens targeted in preclinical PRIT models of solid tumors, the CD138 antigen is moderately expressed (around 10E4 site) per human triple-negative breast cancer cell. Increasing treatment efficacy therefore requires higher doses to be delivered to the tumor in PRIT than in RIT at MTD.

In preparation for a clinical trial of radioimmunotherapy for spontaneous triple-negative canine mammary carcinoma, which is a relevant preclinical model for human triple-negative breast cancer, we established a xenogeneic model using the Mandy cell line, derived from a canine breast tumour biopsy implanted into immunodeficient mice. We had previously isolated monoclonal antibodies specific for the canine CD138 antigen, which is expressed, as in humans, in triple-negative canine mammary cancer ^10^.

PRIT reduces the dose in the blood and increases the therapeutic index compared to RIT. This approach enables the use of higher injected activity without reaching the hematological toxicity limits of conventional RIT. F(ab)’2 fragments exhibit accelerated pharmacokinetics compared to their corresponding IgG, with a shorter time to reach maximum accumulation in the tumor and faster clearance from the blood. These pharmacokinetic properties make F(ab)’2 fragments well-suited for the PRIT approach, particularly when using short-lived isotopes such as the alpha-emitter 211At (half-life: 7.2 hours).

To select a radioimmunotherapy methodology applicable to veterinary clinics, we compared the biodistribution of the canine anti-CD138 antibody 8A1 in its native form and as an F(ab)’2 fragment. To our knowledge, F(ab)’2 antibody fragments have not been used in PRIT approaches employing tetrazine (Tz) and transcyclooctene (TCO) click chemistry for in vivo radiolabeling. We conducted the same analysis on a second tumor model, established using the murine colon adenocarcinoma C15 cell line implanted in immunodeficient mice. The C15 line expresses human CEA at a higher level than CD138 on the Mandy cell line, allowing us to evaluate the influence of tumor antigen expression on the potential efficacy of PRIT using bioorthogonal chemistry with 177Lu or 211At. The human CEA-specific antibody T84.66 (T84) and its F(ab)’2 fragment were used to target CEA in the C15 model.

The biodistribution data were used to evaluate the potential doses delivered to the tumor and to healthy organs in RIT. We also assessed the value of IgG and F(ab)’2 fragments in a PRIT approach using lutetium-177 (^177^Lu) or ^211^At.

This study demonstrated that the F(ab)’2 fragments exhibit comparable pharmacokinetic behavior in these two models and present an advantage over native antibodies for one-step targeting or pre-targeting approaches.

## Results

### Biodistribution of IgG T84 and IgG 8A1 and of their F(ab)’2 fragments

We first evaluated the biodistribution of IgG and F(ab)’2 fragments of the T84.86 (T84) antibody specific for human CEA on the C15 colorectal cancer cell line derived from the MC38 line transfected with human CEA, as well as the 8A1 antibody and its F(ab’)2 fragment specific for the canine CD138 antigen on a triple-negative canine mammary cancer cell line implanted in nude mice. As expected, the F(ab)’2 fragments were cleared much more rapidly from the blood than their parental IgG. After endocytosis by the endothelial cells of blood vessels, IgG capable of interacting with the neonatal Fc fragment receptor (FcRN) are recycled to the vascular compartment, whereas F(ab)’2 are degraded after endocytosis by the lysosomal pathway, leading to a reduction in their biological half-life.

The smaller size of the F(ab)’2 fragments compared to IgG allows for faster penetration into the interstitial milieu of tumors. This is illustrated by higher tumor-to-blood ratios for F(ab)’2 fragments than for IgG, allowing greater vector binding to antigens at the earliest stages of biodistribution for F(ab)’2 fragments. Maximum %ID/g in the tumor was observed 16 hours after injection of the F(ab)’2 fragment compared with 24 hours for IgG T84, and as early as 8 hours for the F(ab)’2 8A1 fragment compared with 24 to 48 hours for IgG 8A1. The biodistribution and organ-to-blood ratio profiles for tumors differ for the IgG and F(ab)’2 T84 vectors compared with IgG and F(ab)’2 8A1. For the former, after a rapid increase in organ-to-blood ratios, a decrease is observed after 24 and 48 hours, respectively, for F(ab)’2 and IgG T84, in contrast to F(ab)’2 and IgG 8A1, for which these ratios increased continuously until the end of the observation period. It is likely that these differences in organ-to-blood ratio profiles reflect differences in the rate of internalization between CEA and CD138. In line with this hypothesis, the tumor-to-blood ratio of a preclinical trial targeting CD138 in a nude mouse model transplanted with the human triple-negative breast cancer cell line MDA-MB-468 shows a progression profile comparable to that observed for the 8A1 antibody in the Mandy line of triple-negative canine breast cancer.

With regard to biodistribution to healthy organs, IgG has a partially constant organ-to-blood ratio over the observation period. This reflects a dynamic equilibrium between vector concentrations in the blood compartment and the interstitial milieu of organs without non-specific binding. The organ-to-blood ratios are proportional to the blood and interstitial volume of healthy organs.

For the F(ab)’2 fragment, the organ-to-blood ratios for the liver and kidney are slightly higher than those for IgG, with little change up to 48 hours after injection. At later time points, these ratios evolve differently for the fragments of the two antibodies. The ratios increase more markedly in the liver with F(ab)’2 8A1 and in the kidney with F(ab)’2 T84 fragments. However, it is notable that at 48 hours, the %ID/g blood for F(ab)’2 8A1 and T84 were only 0.47 ± 0.07 and 1.37% ± 0.09, respectively. These late increases in the organ-to-blood ratio therefore have little influence on the total dose of radioactivity deposited in these healthy organs. To compare the different vectors, we performed a dose calculation for RIT using radiolabeled antibodies to ^131^I.

### Dose assessment for RIT using **^131^I** labelled vector

Vectors directly labeled with iodine on tyrosines are rapidly degraded after internalization, and iodine is excreted by the cells in the form of iodotyrosine. The biodistribution data obtained using an iodine-125 (^125^I) radiolabeled antibody, once corrected for the decay of this isotope, therefore essentially provide information on the quantity of antibody bound to the antigen on the cell surface and present in the blood and interstitial volume of the tumor. For healthy organs that do not express the antigen, the biodistribution values correspond essentially to the antibodies in the blood and interstitial volume. Applying the decay of ^131^I to the biodistribution data obtained with ^125^I enables assessment of the doses potentially deposited in the organs for a RIT performed with ^131^I (Table I).

**Table 1:**
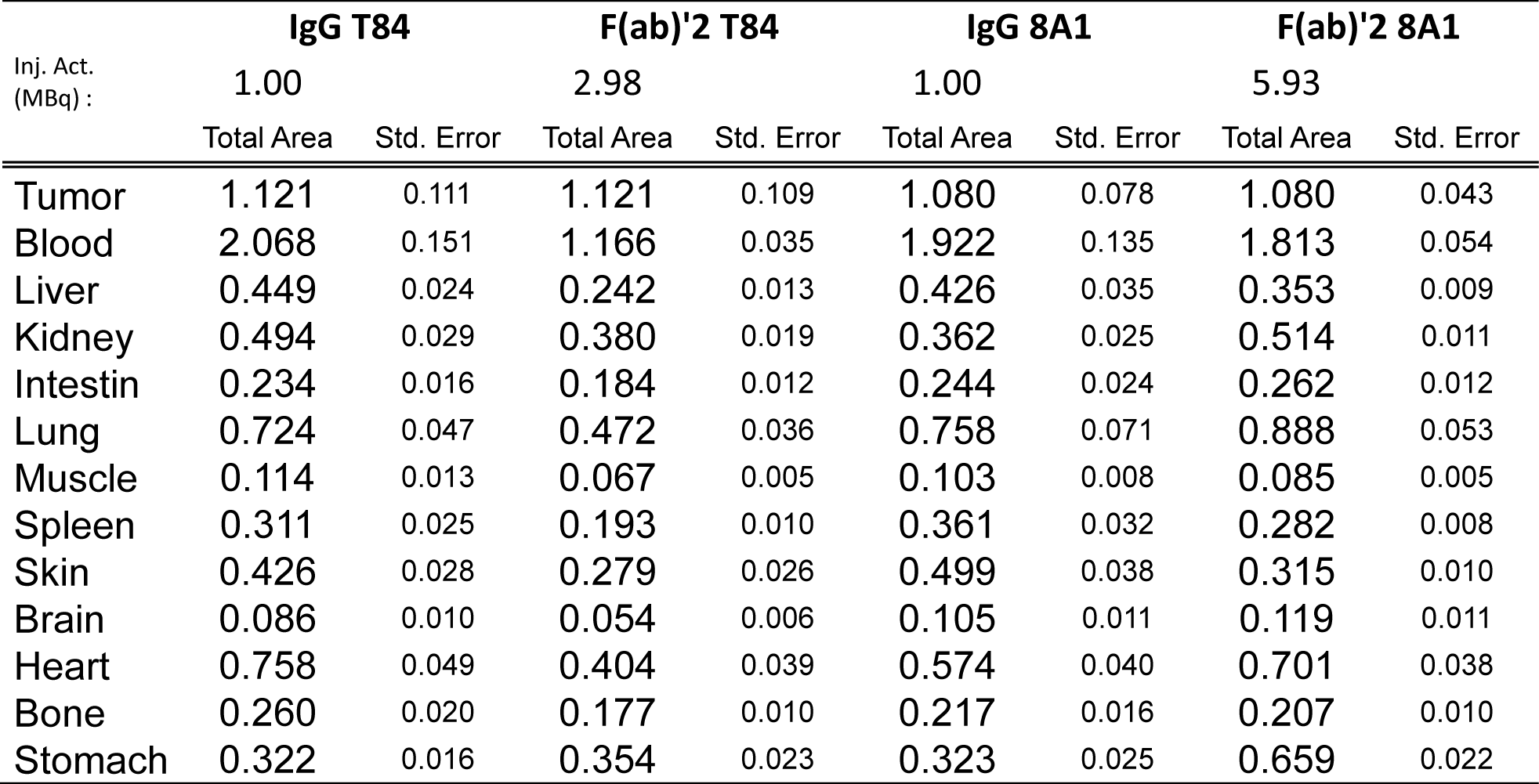
Dose deposited (Gy/MBq injected) to the organs of the C15 tumor targeted by IgGT84.66 or its F(ab)‘2 fragment and to the Mandy tumor by IgG 8A1 or its F(ab)’2 fragment for an ^131^I RIT.

| Inj. Act.<br>(MBq) : | IgG T84 |  | F(ab)'2 T84 |  | IgG 8A1 |  | F(ab)'2 8A1 |  |
| --- | --- | --- | --- | --- | --- | --- | --- | --- |
|  | 1.00 |  | 2.98 |  | 1.00 |  | 5.93 |  |
|  | Total Area | Std. Error | Total Area | Std. Error | Total Area | Std. Error | Total Area | Std. Error |
| Tumor | 1.121 | 0.111 | 1.121 | 0.109 | 1.080 | 0.078 | 1.080 | 0.043 |
| Blood | 2.068 | 0.151 | 1.166 | 0.035 | 1.922 | 0.135 | 1.813 | 0.054 |
| Liver | 0.449 | 0.024 | 0.242 | 0.013 | 0.426 | 0.035 | 0.353 | 0.009 |
| Kidney | 0.494 | 0.029 | 0.380 | 0.019 | 0.362 | 0.025 | 0.514 | 0.011 |
| Intestin | 0.234 | 0.016 | 0.184 | 0.012 | 0.244 | 0.024 | 0.262 | 0.012 |
| Lung | 0.724 | 0.047 | 0.472 | 0.036 | 0.758 | 0.071 | 0.888 | 0.053 |
| Muscle | 0.114 | 0.013 | 0.067 | 0.005 | 0.103 | 0.008 | 0.085 | 0.005 |
| Spleen | 0.311 | 0.025 | 0.193 | 0.010 | 0.361 | 0.032 | 0.282 | 0.008 |
| Skin | 0.426 | 0.028 | 0.279 | 0.026 | 0.499 | 0.038 | 0.315 | 0.010 |
| Brain | 0.086 | 0.010 | 0.054 | 0.006 | 0.105 | 0.011 | 0.119 | 0.011 |
| Heart | 0.758 | 0.049 | 0.404 | 0.039 | 0.574 | 0.040 | 0.701 | 0.038 |
| Bone | 0.260 | 0.020 | 0.177 | 0.010 | 0.217 | 0.016 | 0.207 | 0.010 |
| Stomach | 0.322 | 0.016 | 0.354 | 0.023 | 0.323 | 0.025 | 0.659 | 0.022 |

The doses deposited on tumor C15 by antibody T84 and on tumor Mandy with antibody 8A1 are close, while the maximum %ID/g on the tumor is higher for antibody T84 than for antibody 8A1 (respectively 10.42 %ID/g at 24 h and 7.49 %ID/g at 48 h). The low binding of the 8A1 antibody to the tumor is compensated for by its longer persistence in the tumor compared with the T84 antibody. These differences in binding kinetics are also reflected in lower doses to the tumor with the F(ab)’2 8A1 fragment than with F(ab)’2 T84 (0.182 and 0.376 Gy/MBq).

Besides these differences in doses deposited in the tumor, those evaluated in healthy organs are homogeneous for each kind of IgG or F(ab)’2 vector of the same isotype (murine IgG1). We then studied the dose deposited in healthy organs by IgG and its respective F(ab)’2 fragment for an equivalent dose in the tumor in order to determine which vector format was best suited for RIT.

### Healthy organ dose comparisons for RIT using IgG and F(ab)’2

It is possible to deliver an equivalent dose to the tumor with the same molar quantity of IgG and F(ab)’2 vector by adjusting their specific activity. In our models, the specific activity of F(ab)’2 T84 and 8A1 would be 2.98 and 5.93 times respectively that of the corresponding IgG to reach equivalent dose to tumor (Table 2).

**Table 2:** Comparison of organ dose (Gy) by IgG and F(ab)‘2 for equivalent dose deposited to tumor for ^131^I RIT in C15 (IgG and F(ab)’2 T84.66) and Mandy (IgG and F(ab)’2 8A1) tumor models.

| Gy/MBq | IgG T84 |  | F(ab)'2 T84 |  | IgG 8A1 |  | F(ab)'2 8A1 |  |
| --- | --- | --- | --- | --- | --- | --- | --- | --- |
|  | Total Area | Std. Error | Total Area | Std. Error | Total Area | Std. Error | Total Area | Std. Error |
| Tumor | 1.121 | 0.111 | 0.376 | 0.037 | 1.080 | 0.078 | 0.182 | 0.007 |
| Blood | 2.068 | 0.151 | 0.391 | 0.012 | 1.922 | 0.135 | 0.306 | 0.009 |
| Liver | 0.449 | 0.024 | 0.081 | 0.005 | 0.426 | 0.035 | 0.060 | 0.002 |
| Kidney | 0.494 | 0.029 | 0.128 | 0.006 | 0.362 | 0.025 | 0.087 | 0.002 |
| Intestin | 0.234 | 0.016 | 0.062 | 0.004 | 0.244 | 0.024 | 0.044 | 0.002 |
| Lung | 0.724 | 0.047 | 0.158 | 0.012 | 0.758 | 0.071 | 0.150 | 0.009 |
| Muscle | 0.114 | 0.013 | 0.022 | 0.002 | 0.103 | 0.008 | 0.014 | 0.001 |
| Spleen | 0.311 | 0.025 | 0.065 | 0.003 | 0.361 | 0.032 | 0.048 | 0.001 |
| Skin | 0.426 | 0.028 | 0.094 | 0.009 | 0.499 | 0.038 | 0.053 | 0.002 |
| Brain | 0.086 | 0.010 | 0.018 | 0.002 | 0.105 | 0.011 | 0.020 | 0.002 |
| Heart | 0.758 | 0.049 | 0.136 | 0.013 | 0.574 | 0.040 | 0.118 | 0.006 |
| Bone | 0.260 | 0.020 | 0.059 | 0.003 | 0.217 | 0.016 | 0.035 | 0.002 |
| Stomach | 0.322 | 0.016 | 0.119 | 0.008 | 0.323 | 0.025 | 0.111 | 0.004 |

In regard to IgG T84 and its F(ab)’2 fragment, the doses deposited in healthy organs, and more particularly in the blood, are lower with F(ab)’2 than with IgG for an equivalent dose to the tumor. Less toxicity can thus be expected in the red marrow limiting hematological toxicity. More generally, doses to healthy organs for an equivalent dose to the tumor are lower with the F(ab)’2 T84 fragment than with IgG T84.

No increase in dose to the kidney was observed with injected doses of F(ab)’2 T84 fragments three times higher than those of the corresponding IgG, despite the fact that F(ab)’2 fragments degrade more rapidly than IgG.

In regard with IgG 8A1 and its F(ab)’2 fragment, a specific activity of fragment F(ab)’2 8A1 5.93 times greater than that of IgG T84 is required to achieve an equivalent dose to the tumor. At these injected F(ab)2 fragment activities, the doses to the kidney, lung, heart and stomach are significantly increased compared with IgG but remain lower or practically equivalent for the other healthy organs.

In an earlier preclinical trial, we showed that RIT of the human triple-negative breast cancer line using the CD138-specific B-B4 antibody at the maximum tolerated injected dose corresponding to an injected activity of 22MBq, tumors were eradicated in 50% of the mice over the 100-day follow-up period and their growth severely restricted in the other half of the animals.

Without prejudging the therapeutic effect of the F(ab)’2 T84 and 8A1 antibodies and fragment on the targeted tumors, B-B4 RIT can be used to assess the toxicity of the T84 and 8A1 vectors for an injected activity corresponding to the MTD in the MDA MB 468 targeted with the B-B4-^131^I in nude mice (Table 3)

**Table 3:** Comparison of the dose to organs with IgG and F(ab)’2 T84.66 and 8A1 (Gy) for an RIT with ^131^I at the equivalent dose to the tumor obtained by targeting MDA MB 463 human triple-negative breast cancer with anti-CD138 B-B4 IgG at the MTD.

|  | IgG T84 |  | F(ab)'2 T84 |  | IgG 8A1 |  | F(ab)'2 8A1 |  | BB4 |  |
| --- | --- | --- | --- | --- | --- | --- | --- | --- | --- | --- |
| Act.Inj.<br>(MBq) | 29.9 |  | 89.0 |  | 31.0 |  | 183.9 |  | 22.0 |  |
|  | Total Area | Std. Error | Total Area | Std. Error | Total Area | Std. Error | Total Area | Std. Error | Total Area | Std. Error |
| Tumor | **33.48** | 3.32 | **33.48** | 3.26 | **33.48** | 2.42 | **33.48** | 1.34 | **33.48** | 3.80 |
| Blood | **61.77** | 4.52 | **34.84** | 1.06 | **59.59** | 4.19 | **56.21** | 1.69 | **37.05** | 2.78 |
| Liver | **13.42** | 0.71 | **7.23** | 0.40 | **13.20** | 1.08 | **10.94** | 0.28 | **9.84** | 0.51 |
| Kidney | **14.76** | 0.88 | **11.35** | 0.57 | **11.21** | 0.78 | **15.93** | 0.33 | **8.63** | 0.65 |
| Intestin | **6.99** | 0.47 | **5.51** | 0.37 | **7.55** | 0.74 | **8.14** | 0.36 | **3.26** | 0.23 |
| Lung | **21.63** | 1.40 | **14.08** | 1.07 | **23.51** | 2.20 | **27.54** | 1.65 | **14.99** | 1.21 |
| Muscle | **3.40** | 0.40 | **2.00** | 0.14 | **3.19** | 0.25 | **2.64** | 0.14 | **2.42** | 0.14 |
| Spleen | **9.29** | 0.75 | **5.76** | 0.31 | **11.19** | 1.01 | **8.74** | 0.25 | **7.33** | 0.47 |
| Skin | **12.73** | 0.84 | **8.35** | 0.77 | **15.46** | 1.19 | **9.76** | 0.29 | **9.70** | 0.56 |
| Brain | **2.58** | 0.29 | **1.61** | 0.19 | **3.26** | 0.35 | **3.68** | 0.34 | **1.55** | 0.16 |
| Heart | **22.63** | 1.45 | **12.08** | 1.15 | **17.81** | 1.23 | **21.73** | 1.18 | **11.64** | 0.81 |
| Bone | **7.76** | 0.60 | **5.28** | 0.30 | **6.73** | 0.50 | **6.41** | 0.29 | **5.83** | 0.52 |
| Stomach | **9.62** | 0.48 | **10.57** | 0.70 | **10.03** | 0.77 | **20.43** | 0.68 | **5.01** | 0.28 |

An injected activity of F(ab)’T84 four times higher (89 vs 22MBq) is necessary to achieve a tumor dose equivalent to that obtained in RIT with the B-B4 antibody at the MTD. At this injected activity, the dose to the blood with F(ab)’2 T84 remains lower than that observed with the B-B4 antibody. An identical or slightly lower hematological toxicity is therefore expected. Activities in the liver, lung, muscle, spleen, skin and bone are also slightly lower, while doses in the kidney, intestine and stomach are higher.

Targeting would be of better quality for an RIT performed with the F(ab)’2 T84 fragment than with the corresponding IgG with which blood doses would be 1.7-fold higher than with the MTD of the B-B4 antibody.

For the 8A1 antibody and its F(ab)’2 fragment, the blood doses would represent respectively 1.6 and 1.5 times those obtained with the B-B4 antibody for an equivalent dose to the tumor. The C138 antigen expression on Mandy tumor is thus probably to low to permit efficient RIT with either IgG or F(ab)’2 fragment

This extrapolation of the dosimetry data for RIT using IgGB-B4-^131^I therefore shows that contrarily to IgG 8A1, the F(ab)’2 T84 fragment compatible with animal survival and could be used for RIT. However, the dose to the blood remains higher than that deposited in the tumor, whatever the vector used for RIT in these models expressing the targeted antigen moderately or weakly.

One way of avoiding these high blood doses is to perform a PRIT. Using biodistribution data, we sought to assess whether, as for RIT, F(ab)’2 fragments offered an advantage over IgG for tumor targeting in PRIT approach.

### Evaluation of the potential of IgG and its F(ab)’2 fragment for PRIT

The efficacy of *in-vivo* radiolabeling of tumor-specific vectors in PRIT is low, regardless of the PRIT method used. The dose to the tumor is obtained essentially by the residualization of the isotopes internalized by the tumor. In PRIT assays using click chemistry and ^177^Lu radiolabeled hapten, the dose to the tumor increases with time due to the accumulation of isotopes carried by the radiolabeled vector in the blood. When a clearance agent is used to eliminate circulating vectors before the injection of radiolabeled hapten, this increase in tumor dose is no longer observed.

As mentioned above, biodistribution using ^125^I essentially accounts for isotopes bound to surface antigens and in the blood and interstitial volume of the tumor. It therefore does not provide information on the dose deposited in the organs when the RIT is performed with an internalizing isotope. In this case, the dose no longer depends solely on the quantity of antibodies present in the organs, but also on the rate of internalization of the bound vectors and, in the case of PRIT, on the biodistribution of the radiolabeled hapten and the efficiency of coupling the radiolabeled hapten to the vector pre-localized in the organs and the tumor.

However, the internalization of radiolabeled vectors by the tumor remains directly dependent on the quantity of vector attached to its surface antigen. This data can be measured from the biodistribution carried out with ^125^I. Rossin et al. evaluated the efficiency of *in-vivo* radiolabeling in the TCO/Tz system ^11^. The efficiency of the coupling reaction of radiolabeled Tz to the vector TCO is identical in the tumor and blood. The area under the curve of the biodistribution data for the tumor and organs is therefore independent of the *in-vivo* radiolabeling rate and proportional to the quantity of vector resident in the organs.

By applying the decay of ^177^Lu and ^211^At to the biodistribution data for ^125^I radiolabeled vectors for different injection times of radiolabeled hapten, we calculated the ratios of the areas under the biodistribution curve of healthy organs or tumor to that of blood. The value of these ratios allows comparison of the potential efficacy of a PRIT performed using IgG or its F(ab)’2 fragment, to assess the therapeutic potential of different vectors radiolabeled with ^177^Lu or ^211^At and the toxic risk for healthy organs for different injection schemes (Fig. 2).

**Figure 1:**
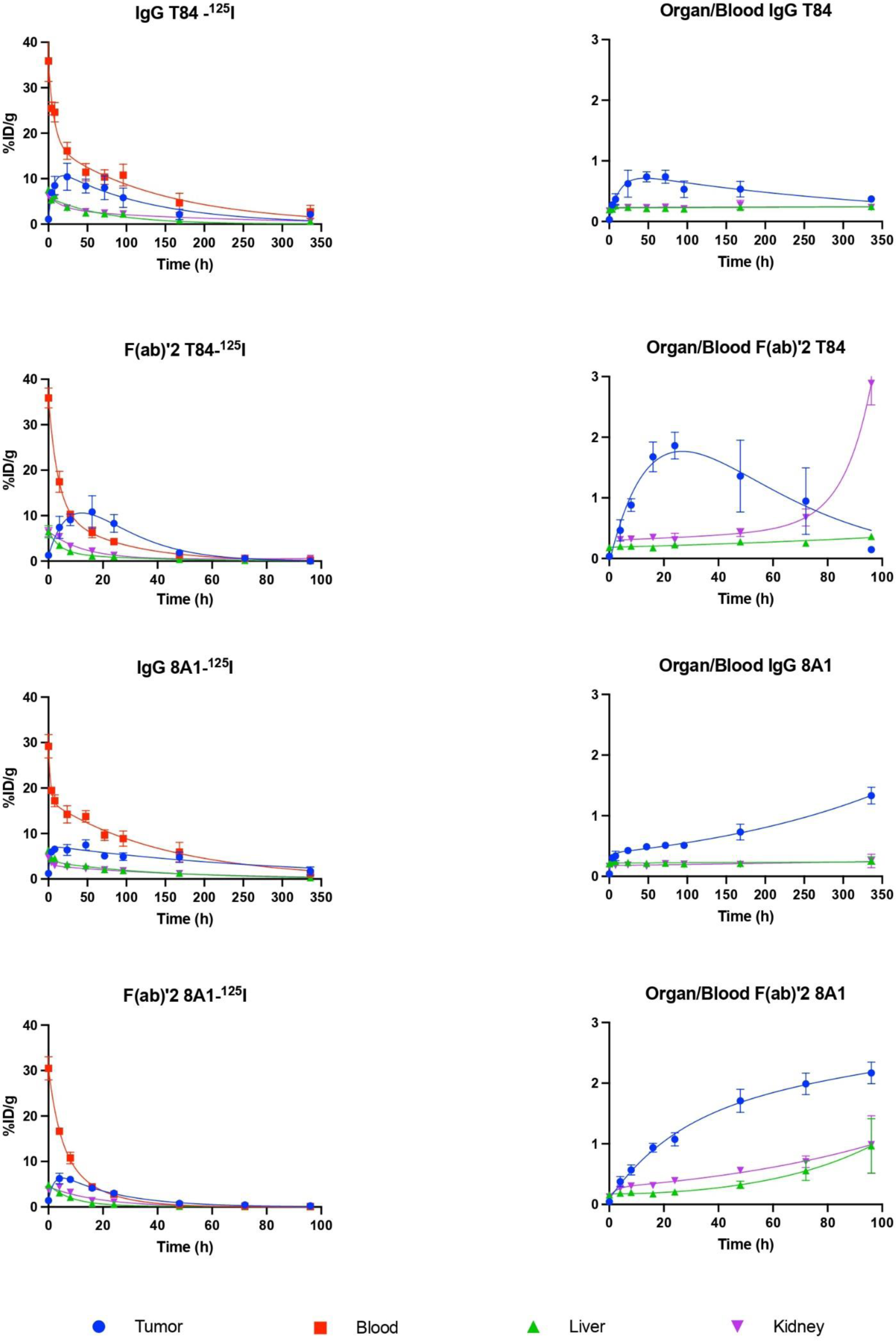
Biodistribution and organ-to-blood ratio of IgG T84 and its F(ab)’2 fragment on C15 tumor and IgG 8A1 and its F(ab)’2 fragment on Mandy tumor tumor. NMRI nude mice engraft with C15 or Mandy tumor cell lines were injected with respectively with T84 IgG and F(ab)’2 fragment or 8A1 IgG and F(ab)’2 fragment radiolabeled with ^I25^I. for each iodinate IgG and F(ab)’2, the biodistribution was evaluated and corrected for 125I decay (left panel) and the organ to blood ratio were calculated fot tumor, blood, kidney and liver at each time point of biodistibution (right panel).

**Figure 2:**
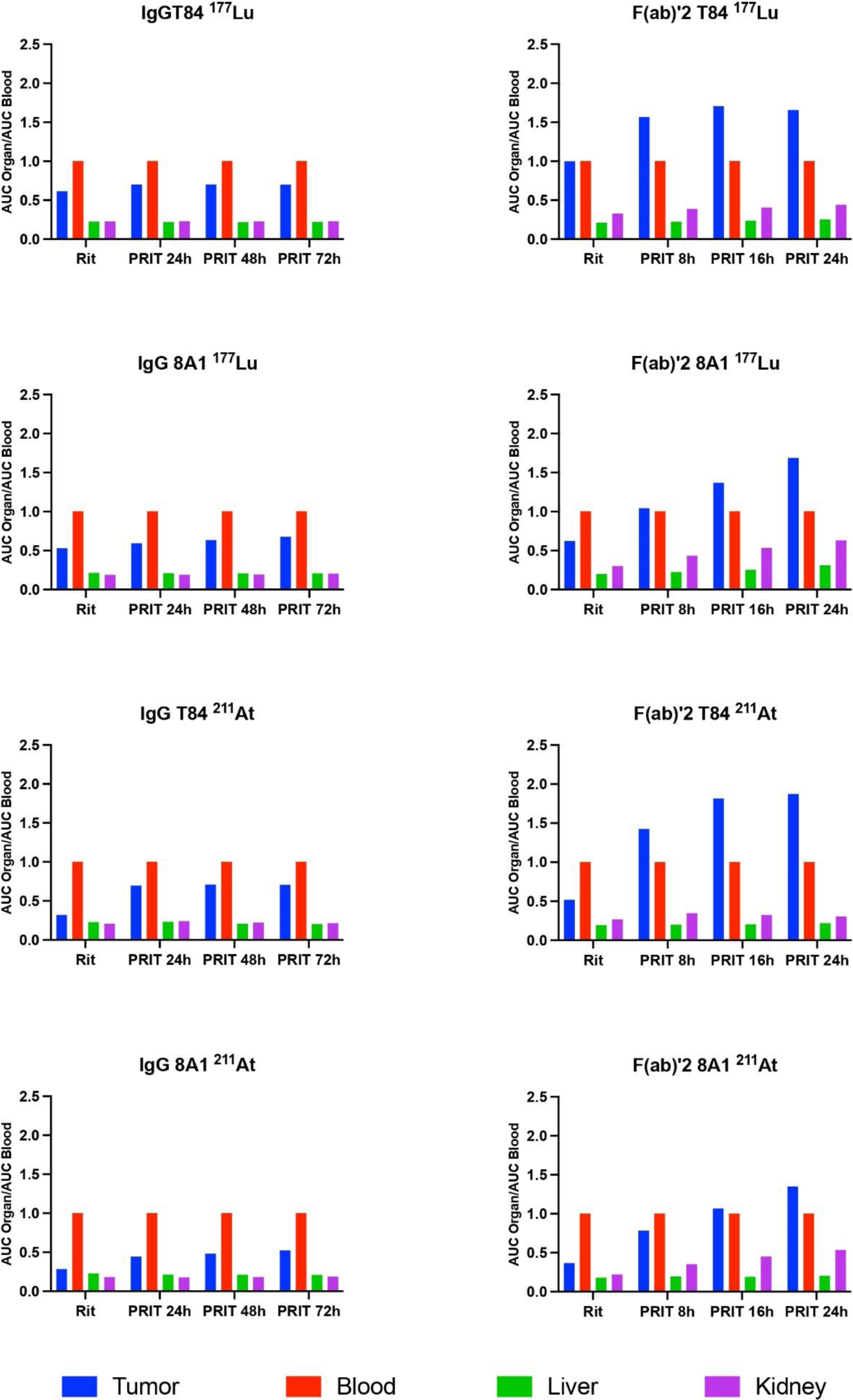
Organ-to blood ratio of the AUC of the curve of in-vivo biodistribution of IgG 8A1-^125^I and its fragment F(ab)’2-^125^I on the Mandy tumor and IgG T84-125I and its F(ab)’2-^125^I fragment on C15 tumor.

### Organ to blood ratio for RIT and ^177^Lu PRIT with IgG and their F(ab)’2 fragments

For RIT and PRIT with ^177^Lu, there was little change in the tumor-to-organ ratio for IgG T84 and 8A1. Whatever the injection time between the vector and the radiolabeled hapten, the tumor-to-blood ratio remained below unity (a maximum of 0.71 for a PRIT at 48h with IgG T84 compared with 0.62 for RIT and a maximum of 0.68 for a PRIT at 72h with IgG 8A1 compared with 0.53 for RIT). The ratios of liver and kidney to blood remained virtually constant regardless of the injection scheme (from 0.22 to 0.23 and 0.23 for liver and kidney respectively with IgG T84 and 0.21 and from 0.19 to 0.20 respectively for liver and kidney with IgG 8A1).

On the other hand, the use of F(ab)’2 fragments led to an improvement in the tumor-to-blood ratio, with a maximum of 1.70 for a PRIT at 16h with F(ab)’2 T84 compared with 1,00 for RIT and a maximum of 1.69 for a PRIT at 24h with F(ab)’2 8A1 compared with 0.63 for RIT. The liver-to-blood ratios are close to those observed for IgG (0.21 and 0.20 for RIT and 0.26 and 0.29 for PRIT at 24h respectively with F(ab)’2 T84 and 8A1). The kidney-to-blood ratios for the F(ab)’2 fragments tended to increase with the delay in injection of the radiolabeled hapten, from 0,33 for RIT to 0.44 for PRIT at 24h for F(ab)’2 T84 and from 0.30 for RIT to 0.63 for PRIT at 24h with F(ab)’2 8A1. These increases in the kidney-to-blood ratio reflect the lower stability of the F(ab)’2 fragment compared with the corresponding IgG.

### Organ to blood ratio for RIT and PRIT at ²¹¹At with IgG and their F(ab)’2 fragments

In contrast to PRIT with ^177^Lu, the use of ^211^At improved tumor-to-organ ratios with IgG and F(ab)’2 fragments. For IgG, the tumor-to-blood ratio increased from 0.32 to 0.71 for RIT and PRIT at 48h with IgG T84 and from 0.28 to 0.52 for RIT and PRIT at 72h with IgG 8A1. Regardless of the injection schedule, the tumor-to-blood ratio was always less than one for the tumors targeted with IgG. This is not the case for the F(ab)’2 fragment, where the ratios became greater than one for a PRIT with an injection delay of 8 hours for F(ab)’2 T84 and 16 hours for F(ab)’2 8A1.

Regarding the liver and kidney, regardless of the injection schedule, the ratios remained constant and comparable to those observed with ^177^Lu (between 0.21 and 0.23 for the liver and between 0.18 and 0.24 for the kidney with IgG T84 and 8A1). For F(ab)’2 fragments, the liver-to-blood ratio changed little regardless of the injection schedule (between 0.19 and 0.18 for RIT and between 0.20 and 0.19 for PRIT at 16h with F(ab)’2 T84 and F(ab)’2 8A1, respectively).

With both ^211^At and ^177^Lu, the kidney-to-blood ratios of F(ab)’2 fragments were higher than those observed with the corresponding IgG and increased with the injection time of the radiolabeled hapten. The ratios increased from 0.27 to 0.35 with F(ab)’2 T84 and from 0.22 to 0.53 with F(ab)’2 8A1 for RIT and PRIT, respectively, for an injection time of the radiolabeled hapten with ^211^At.24h of 8h for F(ab)’2 T84 and 24h for F(ab)’28A1

As mentioned above, these values do not take into account the internalization and residualization of isotopes within tumor cells, which would tend to increase the dose of radioactivity in the tumor and therefore the tumor-to-blood ratio. Healthy organs that do not express the antigen are little affected by antibody internalization. These tumor-to-blood ratios therefore provide qualitative information on the potential toxicity to healthy organs of a PRIT performed using IgG or its F(ab)’2 fragment. However, they provide no information on the availability of the various radiolabeled vectors once they have been distributed in the body, or on the doses that can potentially be delivered to the tumor and healthy organs by any of these vectors.

### IgG and F(ab)’2 fragment residences in tumor and healthy organs for RIT and PRIT

To assess the potential efficacy of IgG and F(ab)’2 fragments in PRIT, we compared the areas under the curve (AUC) of IgG and F(ab)’2 biodistribution data for RIT and PRIT with ²¹¹At or ^177^ Lu for different times of injection of the radiolabeled Tz. As the coupling efficiencies of IgG-TCO to radiolabeled Tz are equivalent in tumor and blood ^11^, we can compare the ratio of the AUC of biodistribution curves to tumor and blood.

In general, the areas under the vector biodistribution curve for RIT or PRIT with ^177^ Lu are much higher for IgG than for F(ab)’2 fragments and decrease in blood and tumor as the injection time between the vector and radiolabeled hapten increases. The AUC value of the tumor biodistribution curve for RIT with IgG T84-^177^ Lu is 3.2 times higher than that for the F(ab)’2 T84-^177^ Lu fragment, and 5.6 times higher for IgG 8A1-^177^ Lu than for the F(ab)’2 8A1-^177^ Lu fragment.

Conversely, the AUC values of the tumor biodistribution data for RIT and PRIT performed using ²¹¹At are of the same order for IgG and its F(ab)’2 fragment. While the AUC values for blood with IgG remain higher than those for tumor regardless of the injection schedule, the AUC values for tumor biodistribution curves with F(ab)’2 fragments are higher than those for blood for a radiolabeled hapten injection delay of 8 hours for F(ab)’2 T84 and 16 hours for F(ab)’2 8A1.

The F(ab)’2 fragments therefore appear to be potentially more suitable for PRIT with ²¹¹At and ^177^ Lu than IgG and allow a greater amount of radiolabeled vector to be delivered to the tumor than to the blood for PRIT. When IgG is used for PRIT, the amount of radiolabeled vector in the tumor remains lower than in the blood, whether ²¹¹At or ^177^ Lu is used in our tumor model of intermediate and low tumoral antigen expression.

Therefore, PRIT makes it possible to improve the tumor-to-blood ratio more markedly with F(ab)’2 fragments than with IgG and with ²¹¹At than with ^177^ Lu. On the other hand, the use of F(ab)’2 fragments reduces the quantities of vector available to the organs, blood, and tumor (Figure 3).

**Figure 3:**
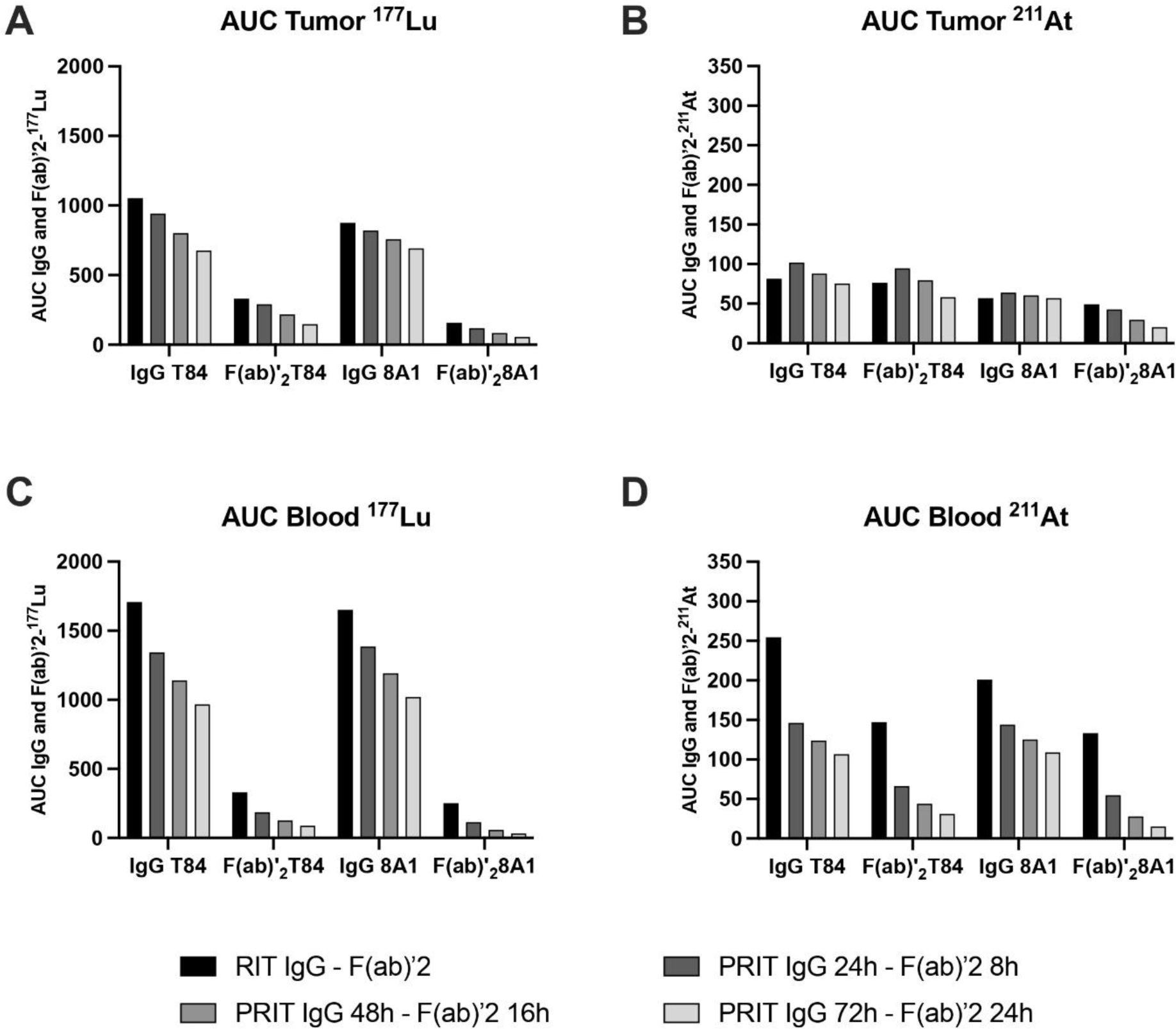
Comparative amount of tumor-available IgG and F(ab)’2 antibodies for RIT and PRIT with different injection times of radiolabeled Tz. The decay of ^177^Lu and ^211^At was applied to the biodistribution curve of IgG and F(ab)’2 T84 and 8A1 (corrected for 125I decay), and the AUC was calculated for each experimental condition. These graphs show the potential amount of radioactive vector in tumor and blood independently of the yield of radiolabeling the vector with Tz-^177^Lu or Tz-^211^At. A and B) AUC of the biodistribution curve in the tumor, C and D) AUC of the biodistribution curve in the blood.

Again, these decreases in the amount of vector available when using the F(ab)’2 fragment instead of IgG were more marked for PRIT using ^177^ Lu than with ²¹¹At.

The lower availability of radiolabeled F(ab)’2 fragments in the tumor compared with IgG can be compensated for by an increase in the specific activity of Tz-^177^ Lu or Tz-²¹¹At, without altering the tumor/blood ratio. The increase in the specific activity (MBq/mol) of radiolabeled Tz required to obtain an equivalent quantity of radiolabeled F(ab)’2 and IgG corresponds to the ratio of their AUC of %ID/g corrected for the decay of ^125^I (% vector injected/g) to which the decay of ^177^ Lu or ²¹¹At from the chosen injection time of radiolabeled Tz is applied.

We compared the increase in AS for a PRIT with IgG for an injection time of 24, 48, and 72h of radiolabeled Tz with the injection time of 8, 16, and 24 h, respectively, of the corresponding F(ab)’2. The AS of radiolabeled Tz required to achieve an equivalent amount of F(ab)’2 to tumor compared with IgG with PRIT is shown in Table 4.

**Table 4:** Ratio of specific activity values required for Tz-^177^Lu or Tz-^211^At to obtain the same AUC value for F(ab)’2 and IgG fragments in the tumors.

| Fold increase radiolabel Tz specific activity | PRIT 8/24 | PRIT 16/48 | PRIT 24/72 |
| --- | --- | --- | --- |
| IgG T84-<sup>177</sup>Lu/F(ab)'2 T84-<sup>177</sup>Lu | 3.24 | 3.70 | 4.56 |
| IgG T84-<sup>211</sup>At/F(ab)'2 T84-<sup>211</sup>At | 1.08 | 1.11 | 1.30 |
| IgG 8A1-<sup>177</sup>Lu/F(ab)'2 8A1-<sup>177</sup>Lu | 6.91 | 9.20 | 12.47 |
| IgG 8A1-<sup>211</sup>At/F(ab)'2 8A1-<sup>177</sup>Lu | 1.50 | 2.04 | 2.80 |

As expected from the comparison of the AUC biodistribution curve, the specific activity of radiolabeled Tz required to achieve tumor-equivalent activities in PRIT with IgG and F(ab)’2 using ²¹¹At is closer than that achieved with ^177^ Lu. Concerning PRIT carried out with ²¹¹At, PRIT at 16 h with F(ab)’2 T84 and 8A1 requires respectively, an injected activity of 1.11 and 2.04 times those necessary for a PRIT with IgG at 48 h to deposit an equivalent activity to the tumor. For PRIT performed with ^177^ Lu, the activity of Tz-^177^ Lu required to attain equivalent activity with F(ab)’2 for a PRIT at 16 h compared to a PRIT with the corresponding IgG at 48 h is respectively 3.70 and 9,20 times the injected activities of IgG T84 and 8A1. We compare the ratio of the blood AUC of IgG radiolabeled with^177^ Lu or ²¹¹At with that of F(ab)’2, for the same quantity of radiolabeled IgG and F(ab)’2 in the tumor by adjusting the specific activity of radiolabeled Tz with the factors determined in Table 4 (Table 5). Regardless of the time of injection of radiolabeled Tz, the quantity of radiolabeled vector in the blood remains lower for PRIT performed with F(ab)’2 than with IgG.

**Table 5:** IgG-to-F(ab)’2 AUC ratio of blood biodistribution curves for equivalent AUCs of tumor biodistribution curves with F(ab)’2 and corresponding IgG (Table 4), for PRIT at different injection times of Tz-^177^Lu or Tz-^211^At.

| AUC blood IgG/AUC blood F(ab)'2 | PRIT 8/24 | PRIT 16/48 | PRIT 24/72 |
| --- | --- | --- | --- |
| IgG T84-<sup>177</sup>Lu/F(ab)'2 T84-<sup>177</sup>Lu | 1.52 | 2.14 | 2.33 |
| IgG T84-<sup>211</sup>At/F(ab)'2 T84-<sup>211</sup>At | 1.96 | 2.04 | 2.37 |
| IgG 8A1-<sup>177</sup>Lu/F(ab)'2 8A1-<sup>177</sup>Lu | 1.12 | 1.63 | 2.01 |
| IgG 8A1-<sup>211</sup>At/F(ab)'2 8A1-<sup>211</sup>At | 1.53 | 1.77 | 2.24 |

### Adaptation of the specific activity of radiolabeled Tz for PRIT compared to RIT with IgG and their F(AB)’2 fragments

The radiolabeling yield of radiolabeled Tz to IgG-TCO in PRIT carried out with ^177^ Lu is usually evaluated with dual isotope biodistribution, where the iodinated vector-TCO provides the %ID/g of accessible vector at the time of injection of radiolabeled Tz, and the %ID/g of the ^177^ Lu radiolabeled vector after conjugation of TCO to Tz-^177^ Lu is estimated 3 hours after the injection of Tz-^177^ Lu. Various variants of Tz and TCO, with their own pharmacokinetics, stabilities *in-vivo*, and efficiency of conjugation, have been evaluated. In some protocols, PRIT was performed with the injection of a clearing agent to remove circulating vectors before the injection of radiolabeled Tz. In these different PRIT protocols, the %ID/g of the vector radiolabeled with Tz-^177^ Lu ranges between 15 and 30% of the %ID/g of the vector-TCO accessible in the tumor ^12 13 14 15^. This means that to obtain equivalent activity in the tumor for PRIT and RIT, the injected activity of radiolabeled Tz must be between 6.66 and 3.33 times that used in RIT. We estimated the increase in the specific activity of radiolabeled Tz required in PRIT carried out with either IgG T84 or 8A1 and their F(ab)’2 fragments to attain the same activity of radiolabeled vectors that would be obtained in RIT with radiolabeled IgG for different efficiencies of radiolabeled Tz conjugation to TCO *in-vivo* (Table 6).

**Table 6:** Fold increase in specific activity of the radiolabeled Tz, for different coupling efficiencies of Tz to TCO, to obtain in PRIT an equivalent AUC of the tumor biodistribution curve of radiolabeled IgG and F(ab)’2 to that obtained with RIT performed with IgG.

| Delay of radiolabeled<br>Tz injection<br>% of radiolabel vector<br>in Tumor | F(ab)'2 8h - IgG 24h |  |  |  | F(ab)'2 16h - IgG 48h |  |  |  |
| --- | --- | --- | --- | --- | --- | --- | --- | --- |
|  | 15% | 20% | 25% | 30% | 15% | 20% | 25% | 30% |
| IgG T84 Lu-177 | 7,4 | 5,6 | 4,5 | 3,7 | 8,8 | 6,6 | 5,3 | 4,4 |
| F(ab)'T84 Lu-177 | **24,2** | **18,1** | **14,5** | **12,1** | **32,5** | **24,3** | **19,5** | **16,2** |
| IgG T84 At-211 | 5,3 | 4,0 | 3,2 | 2,7 | 6,2 | 4,6 | 3,7 | 3,1 |
| F(ab)'2 T84 At-211 | **5,7** | **4,3** | **3,4** | **2,9** | **6,8** | **5,1** | **4,1** | **3,4** |
| IgG 8A1 Lu-177 | 7,1 | 5,3 | 4,3 | 3,6 | 7,7 | 5,8 | 4,6 | 3,9 |
| F(ab)'8A1 Lu-177 | **49,1** | **36,8** | **29,5** | **24,6** | **71,1** | **53,3** | **42,7** | **35,5** |
| IgG 8A1 At-211 | 5,9 | 4,5 | 3,6 | 3,0 | 6,3 | 4,7 | 3,8 | 3,1 |
| F(ab)'2 8A1 At-211 | **8,9** | **6,7** | **5,3** | **4,4** | **12,8** | **9,6** | **7,7** | **6,4** |

In our model of tumors with low and intermediate levels of antigen expression, the Mandy/F(ab)’2 8A1 model requires twice the injected activity of Tz-^177^ Lu or Tz-²¹¹At compared to the C15/F(ab)’2 T84 model. Conversely, for T84 and 8A1 IgG, the increase in the specific activities of Tz-²¹¹At and Tz-^177^Lu required to obtain the same activities in tumor as in RIT are of the same order. The increases in specific activity required for Tz-²¹¹At are less than for Tz-^177^Lu in PRIT with F(ab)’2 and are manageable in terms of radiolabeling. F(ab)’2 fragments therefore offer an advantage over IgG for PRIT with ²¹¹At and, thanks to the increased tumor/blood ratio, make it possible to limit radiation exposure to blood and healthy organs.

## Discussion

### Biodistribution

The biodistribution data for mouse 8A1 and T84 IgG1 isotype antibodies and their F(ab)’2 fragments are very homogeneous for healthy organs. For IgG, the organ-to-blood ratio values were virtually constant over the analysis period (14 days). There is therefore no specific binding to these organs, and the % ID/g observed corresponds to the quantities of antibodies contained at each time in the blood and interstitial volume of the healthy organs.

On the other hand, tumor targeting of CEA on the C15 line with IgG T84 and CD138c on the Mandy line with IgG 8A1 presented different profiles. With IgG T84, the tumor-to-blood ratio decreased from 72 hours after injection, whereas it increased continuously with the 8A1 antibody until the end of the study (14 days). IgG 8A1 persists longer on the tumor surface than IgG T84, although the latter has greater maximal binding to the tumor.

These differences in pharmacokinetic behavior could be linked to the internalization processes of ACE and CD138 antigens. The internalization of ACE has been described as a clathrin-dependent process ^16^, unlike the internalization of CD138, which uses a non-clathrin-coated pit-dependent internalization pathway ^17 18^. Clathrin-coated pit-dependent internalization is much more rapid than clathrin-coated pit-independent internalization. The iodinated vectors we used to assess biodistribution are rapidly degraded once internalized, and the iodine is excreted as iodotyrosine. Slow internalization kinetics could allow for longer persistence of the vector bound to surface antigens such as CD138 compared to the more rapidly internalized and degraded CEA-bound vectors.

### Interest of F(ab)’2 fragments for RIT

The biodistribution profiles and tumor-to-blood ratios of the F(ab)’2 fragments are consistent with those of their respective IgG. Because of their smaller size compared with IgG, F(ab)’2 fragments diffuse more rapidly within the tumor and healthy organs, as evidenced by their higher tumor-to-blood ratios. The kidney-to-blood ratios increase progressively with time in accordance with the elimination of F(ab)’2 fragment degradation products by glomerular filtration. The kidney-to-blood ratio increases more sharply from 48 hours after injection when the quantity of F(ab)’2 fragment has largely been eliminated from the body.

Although the quantities of F(ab)’2 present in the tumor are lower than those of IgG for equivalent molar quantities injected, as shown by the areas under the biodistribution curve (Table 1), F(ab)’2 fragments are clearly of interest for RIT. The dose to blood and healthy organs is reduced in RIT carried out with F(ab)’2 compared to RIT with IgG at equivalent doses to the tumor (Table 2). This is more pronounced for the C15 tumor targeted with F(ab)’2 T84 than for the MDA-MB-468 model expressing lower antigen density on the tumor (Figure 1).

In order to compare the toxicity to healthy organs in RIT performed with F(ab)’2 and IgG, we use the MTD determined in the xenogeneic MDA-MB-468 human triple-negative breast cancer model as a reference ^19^. The lower amount of F(ab)’2 fragment in the tumor compared with IgG can be compensated for by increasing the injected activity using a higher specific activity of the radiolabeled F(ab)’2 vector without altering its biodistribution (Table 3). For a tumor dose equivalent to that in the MDA-MB-468 model, at an injected activity corresponding to the MTD, the use of the F(ab)’2 T84 fragment reduces the dose to the kidneys and blood for an injected activity three times greater than that of IgG (Table 3). These results are consistent with a preclinical study comparing iodinated IgG, F(ab)’2, and F(ab) fragments, in which the MTD of the F(ab)’2 and F(ab) fragments was 4.6 and 11.5 times, respectively, the MTD of the corresponding IgG ^20^. On the Mandy model, 5.9 times the injected activity of the 8A1 IgG would be require to reach tumor-equivalent with the 8A1 F(ab)’2 fragment (Table 2), resulting in blood irradiation greater than the MTD (Table 3). The expression of canine CD138 on the Mandy line is therefore insufficient to allow effective radioimmunotherapy using an ^131^I -labeled antibody due to the high blood irradiation. However, the doses deposited in healthy organs remain of the same order for IgG 8A1 and F(ab)’2 8A1, with the exception of the stomach and kidneys.

On the C15 tumor model, an equivalent dose to the tumor, comparable to that of the B-B4 antibody at the MTD, could be achieved with the F(ab)’2 T84 fragment, resulting in a lower dose to blood, whereas the use of IgG T84 exceeds the MTD and the dose to healthy organs calculated for F(ab)’2 T84. This indicates that the use of the F(ab)’2 fragment in place of IgG could be advantageous for RIT.

Based on these biodistribution data, we aimed to assess whether the use of the F(ab)’2 fragment offers an advantage over IgG for PRIT using click chemistry with the IEDDA functions TCO and Tz.

### Interest of F(ab)’2 fragments for PRIT

We compared the organ and tumor residence of vectors radiolabeled with ²¹¹At t, an alpha emitter with a half-life of 7.2 h, and with ^177^ Lu, a beta emitter with a half-life of 6.7 d. To this end, the decay of ²¹¹At or ^177^ Lu was applied to the biodistribution data for the different vectors radiolabeled with ^125^I, corrected for ^125^I decay, which is representative of the quantities of vector present in the tumor and organs. The areas under the curve were calculated, taking as initial points the different injection times between vector and radiolabeled Tz.

For the F(ab)’2 fragment, the area under the curve was evaluated for injection delays of 8, 16, and 24 hours and 24, 48, and 72 hours for IgG. The AUC of the biodistribution curves can be used to determine, for healthy organs, the quantity of vector available in the blood and interstitial volume. For tumors, these ratios also include the quantity of vector bound to tumor antigens on the surface of tumor cells.

The quantities of IgG available to the tumor in the T84/C15 and 8A1/Mandy models for PRIT with ^177^ Lu are always lower than the quantities of circulating vectors, and the benefit of PRIT compared with RIT would therefore remain very limited. In the two models investigated, the expression of tumor antigens seems too low to allow the accumulation of higher quantities of radiolabeled IgG in the tumor than in the blood. PRIT with ²¹¹At does, however, significantly increase tumor IgG levels relative to blood.

Biodistribution with ^125^I radiolabeled vectors does not take into account the residualization of the metal isotope, such as ^177^ Lu, which increases the dose deposited in the tumor. Specific biodistributions using ^177^ Lu radiolabeled vectors are needed to draw a definitive conclusion on the efficacy of PRIT and RIT using this isotope. However, the amount of internalized radiolabeled vector and residualized isotopes is directly dependent on the amount of vector binding to the tumor cell surface, as determined here with iodinated vectors, and this parameter allows comparison of the potential efficiency of IgG and its F(ab)’2 fragment in PRIT with residualizing isotopes.

With regard to the F(ab)’2 fragments, the quantities of vectors in the tumor are always less than or equal to those in the blood for RIT, whereas they are systematically greater for PRIT (with the exception of F(ab)’2 8A1 for a Tz injection delay of 8 h), regardless of the injection time of the radiolabeled Tz. The F(ab)’2 fragments therefore appear to be better adapted to PRIT than the corresponding IgG. It is noteworthy that the quantities of vector in the tumor for RIT with ²¹¹At using F(ab)’2 fragments are of the same order as those evaluated for IgG (Figure 3).

It therefore seems possible to obtain superior tumor irradiation with F(ab)’2 fragments than with IgG in PRIT by adjusting the specific activities of radiolabeled Tz, without reaching blood doses leading to limiting hematological toxicity. Treatment at the maximum tolerated dose would allow greater tumor irradiation in PRIT with ²¹¹At using an F(ab)’2 fragment instead of IgG for tumor cells expressing low quantities of antigens.

### Adjustment of injected activity of radiolabeled Tz in PRIT

Contrary to SPECT-CT or PET-CT imaging, where the improvement of the contrast between tumor sites and healthy organs with pretargeting is of great interest for diagnosis, the dose to the tumor remains the primary concern for therapy.

The different assays from the literature report that, in the case of PRIT using the TCO/Tz click chemistry, a small proportion of the radiolabeled Tz penetrates the tumor, and that the radiolabeling efficiency is limited in time and quantity. It thus appears that the efficiency of coupling radiolabeled Tz to the TCO of prelocalized vectors is a central parameter to optimize targeting efficiency in PRIT. To address this challenge, optimized variants of TCO ^21^ and Tz ^22^ have been developed to increase *in-vivo* stability and reactivity for the IEDDA reaction. Membreno et al. have developed dendrimers decorated with TCO in order to increase the yield of radiolabeling of tumor-bound IgG-TCO. Despite these intensive efforts, the low amount of radiolabeled Tz penetrating the tumor limits the % ID/g of Tz-^177^ Lu to the range of 15 to 30% of the actual % ID/g of IgG-TCO in the tumor. The use of clearing agents to remove circulating IgG-TCO from the blood in order to increase the tumor-to-organ ratio has also been extensively investigated.

Membreno et al., comparing the biodistribution of the huA33 antibody conjugated to TCO (3 TCO/Ac on average) in a PRIT assay at 72 hours with or without a clearing agent, show that at 4 hours after injection of Lu-DOTA-PEG7-Tz, the % ID/g to the tumor is identical (3.63 and 3.32 with or without a clearing agent) and decreases over time in the presence of a clearing agent, while increasing progressively in its absence (1.93 and 12.66 % ID/g at 120 hours, respectively) ^21^. Most of the dose to the tumor is therefore acquired over time by endocytosis of the radiolabeled vectors and residualization of the isotopes within the tumor cells. This process requires the presence of circulating radiolabeled antibodies that diffuse towards the tumor to fuel the progressive sequestration of the isotopes in the intracellular volume and the increase in dose over time. Tumor % ID/g is much higher using the same antibody in RIT than in PRIT without a clearing agent. Membreno et al., targeting the A33 antigen on the SW1222 tumor, observed a % ID/g of 11% ± 2.4 five days after Tz-^177^ Lu injection for a PRIT with an interval of 48 hours, and 75% ± 22 % ID/g five days after the injection of directly labeled huA33 IgG in RIT. At the same time, the tumor-to-organ ratios were comparable for RIT and PRIT, and the authors concluded from dosimetric analysis that PRIT offers a modest improvement compared to RIT ^7^.

Here we propose to use the F(ab)’2 fragment in place of IgG in two tumor models with intermediate (C15 tumor) to low (Mandy tumor) tumor antigen expression. In RIT, tumor-to-blood ratios are improved with F(ab)’2 compared to IgG when injected activities are adjusted to obtain the same dose to the tumor (Table 2). The therapeutic index was also improved with F(ab)’2 T84 compared to IgG T84 in healthy organs in the C15 tumor and was equivalent for the kidney with both vectors. However, the low tumor antigen expression on the Mandy tumor precludes the application of RIT since the injected activities needed to deposit a therapeutic dose to the tumor would lead to strong irradiation of healthy organs. Conversely, PRIT carried out with F(ab)’2 improves the organ-to-blood ratio compared to IgG. This improvement is more pronounced when ²¹¹At t is used instead of ^177^Lu.

Using biodistribution data from IgG and iodinated F(ab)’2 fragments, we evaluated the amount of vector present in the tumor for different injection times between the vector and Tz-^177^ Lu or Tz-²¹¹At. Tumor-to-blood ratios are improved with F(ab)’2 fragments for a PRIT with²¹¹At and to a lesser extent with ^177^ Lu (Figure 2). With the latter isotope, the doses deposited in the tumor are greatly reduced compared with a PRIT performed using IgG. With ²¹¹At, on the other hand, the quantities of F(ab)’2 fragments in the tumor, although slightly lower, are of the same order as with IgG (Figure 3). We have considered the possibility of compensating for the reduced deposition of the F(ab)’2 fragment in the tumor by adjusting the specific activity of radiolabeled Tz with ^177^ Lu and ²¹¹At for different coupling efficiencies of radiolabeled Tz to IgG-TCO, estimated with dual isotope biodistribution reported in the literature (Table 6). This approach makes it possible to increase the doses deposited in the tumor and in healthy organs while maintaining the biodistribution profile and the tumor-to-blood ratio. The increases in specific activity remain manageable in terms of radiolabeling Tz with ²¹¹At (2.9 and 5.70 times the specific activity required to achieve PRIT with F(ab)’2 the activity to the tumor obtain in RIT with). However, with ^177^ Lu, the specific activities required to achieve a dose equivalent to the tumor in the RIT protocol with the corresponding IgG seem difficult to envisage.

To our knowledge, F(ab)’2 fragments have not yet been evaluated as vectors for PRIT using the IEDDA reaction. This study suggests that they may be of interest to limit the dose to blood compared to IgG for PRIT using isotopes with short half-lives like ²¹¹At. Regarding PRIT using ^177^ Lu, further investigations are needed to evaluate the residualization of this metallic isotope within tumors. In RIT and PRIT, most of the dose to the tumor is acquired over time by endocytosis of the radiolabeled vectors and residualization of the isotopes within the tumor cells. This process requires the presence of circulating radiolabeled antibodies that diffuse towards the tumor to facilitate the progressive sequestration of the isotopes in the intracellular volume and the increase in dose over time, precluding the use of clearing agents for PRIT, at least in the low antigen-expressing tumors investigated in this article. The doses to blood and healthy organs that don’t express the targeted antigens are less affected by internalization than tumors, offering the possibility of an enhanced therapeutic index. The rate of internalization of antibodies by tumor cells is therefore decisive in defining the injected activity required to obtain therapeutic efficacy with acceptable toxicity to healthy organs.

## Material and Method

### Animals

Female NMRI-nu (nu/nu) mice, 7 to 8 weeks old and weighing between 28 and 34 g, were obtained from Janvier LABS (Le Genest-Saint-Isle, France). They were housed under standard conditions in the UTE animal facility (SFR François Bonamy, IRS-UN, Université de Nantes, license number B-44-278). The Pays de la Loire animal experimentation ethics committee validated all the experiments carried out (accreditation No. CEEA 2012-171). For biodistribution assays, mice were transplanted with subcutaneous injections of 2 × 10^6 C15 cells or 5 × 10^6 Mandy cells. Mice were used for biodistribution assays 2 and 6 weeks after the implantation of C15 and Mandy cells, respectively.

### Cell lines

The C15 cell line is derived from the murine MC38 colonic adenocarcinoma cell line by stable transfection of the cDNA encoding human CEA. The Mandy canine mammary cancer line was obtained from ONIRIS (Nantes Veterinary School) and was isolated from a mammary tumor biopsy. The Mandy and C15 cell lines were cultured in RPMI 1640 medium (Sigma Aldrich, Saint-Quentin-Fallavier, France) supplemented with 10% fetal calf serum (SVF) (Fisher Scientific SAS, Illkirch Cedex, France; reference 15313701), L-Glutamine 200 mM (Thermo Fisher Scientific, Illkirch-Graffenstaden, France), and 100 U/mL of penicillin-streptomycin (10,000 U/mL) from the same supplier. The cell lines were grown at 37°C, 5% CO₂, and 100% humidity.

### Antibody

Antibody T84.66 (T84) is a murine IgG1 specific for the human CEA antigen ^23^.Antibody 8A1 is a murine IgG1 specific for the canine CD138 antigen, isolated in the laboratory by immunization of mice with a soluble recombinant form of canine CD138 ^10^.

Hybridomas producing T84 and 8A1 antibodies were cultured in RPMI 1640 media supplemented with 10% IgG-depleted fetal bovine serum in 1 L Erlenmeyer flasks (Corning®, New York, USA) with stirring at 80 rpm for 4 days at 37°C and 5% CO₂ in an agitator. Supernatants from clone cultures were collected, and canine CD138-specific antibodies were purified over a HiTrap Protein G column (GE Healthcare, Piscataway, NJ, USA). Briefly, supernatants from hybridoma cultures were diluted in phosphate buffer (1:1) to adjust the pH to 7. After passage through the column, antibodies were eluted using a glycine–HCl buffer at pH 2.7 and dialyzed overnight against PBS at pH 7.4 using a 30,000 MWCO dialysis cassette (Thermo Scientific, Waltham, USA). Purified antibodies were filtered through 0.2 μm filters and stored at 4°C, and their production yields were determined. Isotypes and light chains of purified antibodies were characterized using a mouse monoclonal antibody isotyping kit (Roche) according to the kit instructions.

### Production of F(ab)’2 fragments

Production of the F(ab’)2 fragment of the 8A1 and T84 antibodies was carried out according to a protocol previously established in the laboratory. 5 mg of 8A1 antibody was added to porcine pepsin (Sigma Aldrich, Saint-Quentin-Fallavier, France) at a pepsin/antibody ratio of 40, in 0.1 M citrate buffer, pH 3.75, for two hours at 37°C. The reaction is stopped by adding 1 M Tris base. After checking the digestion by SDS-PAGE, the F(ab’)2 fragments are purified by FPLC on a Superdex 200 column (GE Healthcare) at a flow rate of 1 ml/min. Fractions containing the F(ab’)2 fragment are then pooled and concentrated on Centricon (Millipore, Sigma Aldrich, Saint-Quentin-Fallavier, France) in PBS.

### Radiolabeling of IgG and F(ab)’2 fragment

Radiolabeling of IgG and F(ab)’2 fragments with ¹²⁵I was carried out according to the iodogenic method. Vector iodination was performed using Iodo-Gen (1,3,4,6-Tetrachloro-3a,6a-Diphenylglycouril, Sigma, Saint-Quentin-Fallavier, France) as the oxidant with ¹²⁵I (NaI, Perkin Elmer, Billerica, MA). Iodo-Gen was dissolved in 0.1 mg/mL dichloromethane, divided into 100 μL fractions in glass tubes, evaporated, and stored at 4°C. The vector and ¹²⁵I were incubated in these tubes for 10 minutes at room temperature with gentle agitation. The Iodo-Gen converts I⁻ ions to I⁺, promoting their binding to the vector tyrosines by an electrophilic substitution reaction. The reaction is stopped by transferring the mixture to a tube without Iodo-Gen.

ITLC™-SG thin-layer chromatography (Pall Corporation, Fontenay-sous-Bois, France) was performed to assess the labeling yield: 1 μL of the labeled solution was deposited at the bottom of the ITLC™-SG plate (9 cm), 1 cm from the bottom. The plate, immersed in 1 mL of 10% trichloroacetic acid (Prolabo, Paris, France), separates the free iodine, which migrates to the top of the plate, from the radiolabeled antibody, which remains at the bottom. After cutting, the bottom of the plate containing the radiolabeled antibody and the top containing the free iodine are counted with a γ counter (1480 Automatic Gamma Counter, Wallac, Billerica, MA), and the labeling yield is calculated.

The radiolabeled vector is purified on a Sephadex PD10 column (Amersham Pharmacia Biotech) with PBS elution. ITLC™-SG chromatography is used to estimate the radiochemical purity of the iodinated vector. The radiochemical purity of the iodinated vectors is greater than 95% for the complete 8A1 antibody and its F(ab’)2 fragment.

### Biodistribution assay

The biodistributions of F(ab’)2 fragments and 8A1 IgG were performed on groups of three mice. Animals were injected into the tail vein with 6 μg of IgG-¹²⁵I or 4 μg of F(ab’)2-¹²⁵I fragments. At the chosen time, the mice were sacrificed, and the organs were collected. After weighing, the radioactivity of the organs was measured in a beta counter. The results are expressed as a percentage of activity per gram of organ (%ID/g). Injected activity was measured at the time of injection by counting, in a beta counter, three doses prepared identically to those injected into the animals. For F(ab’)2 fragments, sacrifice times after injection were 2 min, 4 h, 8 h, 16 h, 24 h, 48 h, 72 h, and 96 h. For IgG, sacrifice times were 2 min, 4 h, 8 h, 24 h, 48 h, 72 h, 96 h, 168 h, and 336 h.

### Analysis of biodistribution

The ¹²⁵I biodistribution data, expressed as %ID/g of organ, were corrected for ¹²⁵I decay to give the amount of carrier present in the organs at each time point. For RIT dosimetry using ¹³¹I, the decay of this isotope was applied to the biodistribution values corrected for ¹²⁵I decay. The number of decays at each organ for an injected activity of 1 MBq was determined at each analysis time, and the corresponding energy was calculated as the product of the number of decays and the mean energy of an ¹³¹I decay. The area under the curve of the values thus obtained is used to determine the self-dose to each organ in Gy.

To calculate the organ-to-blood ratio in PRIT for the F(ab’)2 and IgG 8A1 and T84 fragments, a non-linear “sum of exponentials” regression was performed for the kidney, liver, blood, and tumor using biodistribution data corrected for ¹²⁵I decay (GraphPad PRISM). A curve was generated from the equation constants. The decay of ²¹¹At or ¹⁷⁷Lu was applied to the values of the curve generated from the different times of injection of radiolabeled Tz with ²¹¹At or ¹⁷⁷Lu. The area under the curve was calculated for each injection time of radiolabeled Tz. Organ-to-blood ratios were calculated from the AUC values for each time point. The calculated AUC values were used to compare the amounts of radiolabeled ²¹¹At and vectors potentially present in the tumor and blood for each considered time of injection of radiolabeled Tz.

