## Supplementary figures and images for "Advantage of the F(ab)’2 fragment over IgG for RIT and PRIT"

### Supplemental Data 1

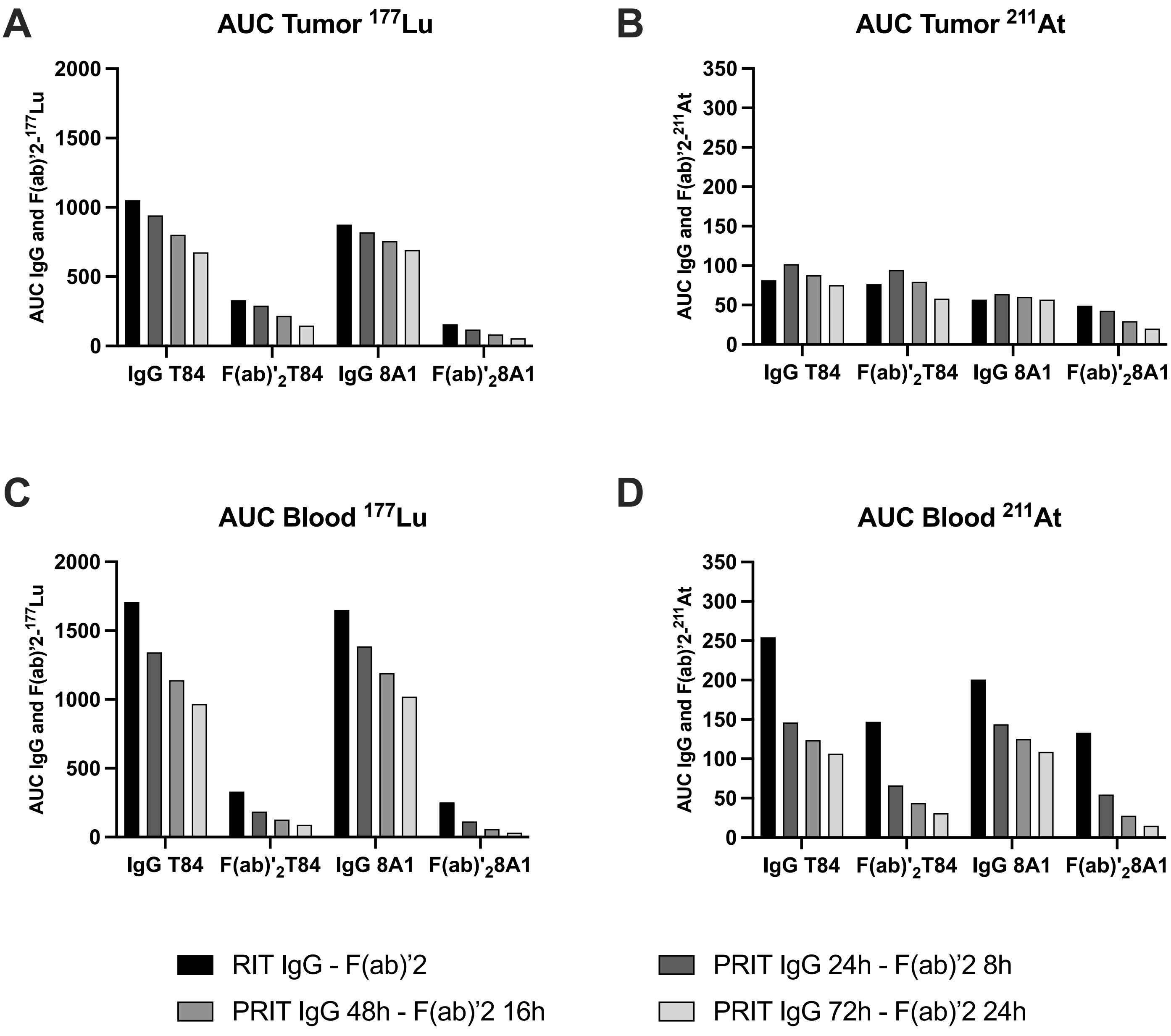
